# A far-red fluorescent chemogenetic reporter for in vivo molecular imaging

**DOI:** 10.1101/2020.04.04.022145

**Authors:** Chenge Li, Alison G. Tebo, Marion Thauvin, Marie-Aude Plamont, Michel Volovitch, Xavier Morin, Sophie Vriz, Arnaud Gautier

## Abstract

Far-red emitting fluorescent labels are highly desirable for spectral multiplexing and deep tissue imaging. Here, we describe the generation of frFAST (far-red Fluorescence Activating and absorption Shifting Tag), a 14-kDa monomeric protein that forms a bright far-red fluorescent assembly with (4-hydroxy-3-methoxy-phenyl)allylidene rhodanine (HPAR-3OM). As HPAR-3OM is essentially non-fluorescent in solution and in cells, frFAST can be imaged with high contrast in presence of free HPAR-3OM, which allowed the rapid and efficient imaging of frFAST fusions in live cells, zebrafish embryo/larvae and chicken embryo. Beyond enabling genetic encoding of far-red fluorescence, frFAST allowed the design of a far-red chemogenetic reporter of protein-protein interactions, demonstrating its great potential for the design of innovative far-red emitting biosensors.

## INTRODUCTION

The high demand for far-red fluorescent labels has emerged from the need for new colors as the growing use of optical biosensors and optogenetic tools obstruct the spectral window available for biological imaging. Moreover, far-red fluorescent labels allow imaging deeper in tissues as the autofluorescence, light scattering and absorbance by endogenous molecules is reduced in the far-red region. The first far-red fluorescent proteins were engineered from phytochromes and phycobiliproteins^1–3 4–6^. These engineered fluorescent proteins covalently bind biliverdin, which fluoresces in the far-red region once attached to the protein. Far-red fluorescent proteins were successfully imaged in cells and animals; however, the availability of intracellular biliverdin and the rather slow rate of chromophore attachment (half-time of maturation > 100 min) was shown to affect the efficiency of labeling^7^.

Chemogenetic fluorescent reporters based on self-labeling tags and fluorogen-activating proteins have been proposed as alternative to encode far-red fluorescence and label proteins in cells and multicellular organisms applying synthetic fluorogenic dyes exogenously^8–11^. Here, we describe frFAST, a small monomeric protein tag of 14 kDa that forms a far-red fluorescent assembly with (4-hydroxy-3-methoxy-phenyl)allylidene rhodanine (HPAR-3OM) (**Figure 1a**) and enables the observation of fusion proteins in live cells and organisms. frFAST is a variant of FAST (Fluorescence-Activating and absorption-Shifting Tag), a monomeric protein tag engineered from the apo photoactive yellow protein (PYP) from *Halorhodospira halophila*. In its original form, FAST forms with an exquisite selectivity fluorescent complexes with derivatives of 4-hydroxybenzylidene rhodanine (HBR)^12,13^ (**Supplementary Figure 1**), which are otherwise non-fluorescent chromophores. This strong fluorogenic effect allows the imaging of FAST-tagged proteins in cells and multicellular organisms with high contrast. Labeling of FAST-tagged proteins in cells takes place within a few seconds and can be reversed by simple washing, if needed. With a size of 14 kDa, FAST is one of the smallest genetically encoded tags, limiting the risk of dysfunctional fusions and minimizing its genetic footprint. In addition, as it does not require molecular oxygen to be fluorescent, FAST was shown to be an attractive alternative to classical fluorescent proteins for imaging biological phenomena in anaerobic systems^14,15^. Introduction of a single mutation (V107I) and/or tandemerization was furthermore shown to be an efficient way to improved brightness^16^. Imaging of proteins below the diffraction limit using super-resolution imaging by radial fluctuations^17^ or single-molecule localization microscopy (SMLM)^18^ could moreover be achieved thanks to the ability to label only a subset of FAST-tagged proteins by adjusting the chromophore concentration. Because of its modular nature, FAST allowed also the design of a Ca^2+^ biosensor^19^ and a fluorescent split reporter, splitFAST, with rapid and reversible complementation for the detection of transient protein-protein interactions^20^.

**Figure 1.**
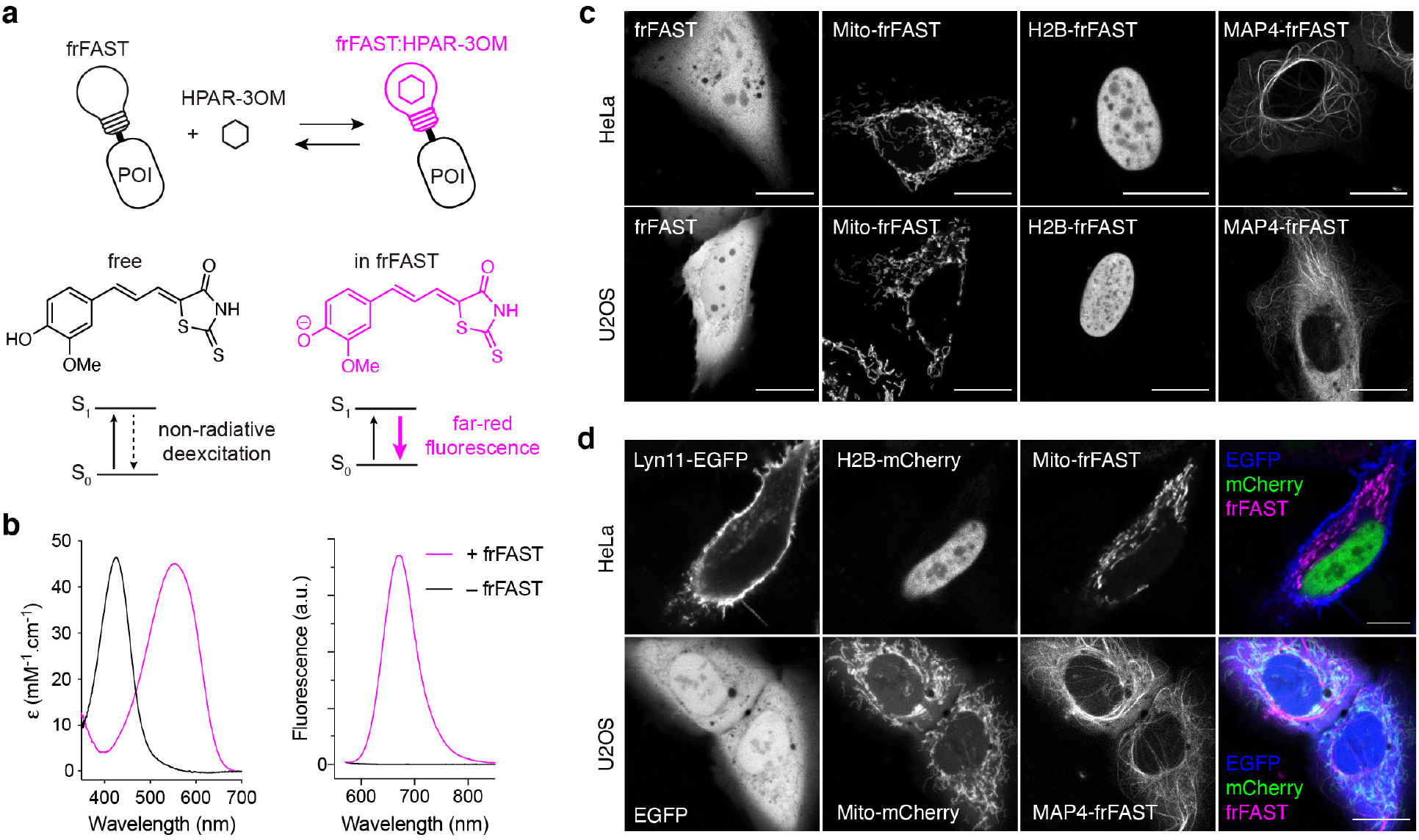
frFAST forms a far-red fluorescent assembly with HPAR-3OM and enables to encode far-red fluorescence in cells. **(a)** frFAST principle and structure of HPAR-3OM (POI: protein of interest). **(b)** Absorption and emission spectra of the HPAR-3OM in absence (black) or presence (magenta) of frFAST. HPAR-3OM and frFAST concentrations were 2 μM and 40 μM, respectively, in pH 7.4 PBS (50 mM sodium phosphate, 150 mM NaCl). Spectra were recorded at 25°C. **(c)** Confocal micrographs of live HeLa and U2OS cells expressing various frFAST fusions. Cells were incubated with 10 μM HPAR-3OM for 15-30 seconds and then directly imaged using the following settings: Ex/Em 633/638-797 nm. **(d)** Confocal micrographs of live HeLa and U2OS cells co-expressing various EGFP, mCherry and frFAST fusions. Cells were incubated with 10 μM HPAR-3OM for 15-30 seconds and then directly imaged using the following settings: EGFP Ex/Em 488/493-599 nm, mCherry Ex/Em 543/560-598, frFAST Ex/Em 633/650-797 nm. **(c,d)** Mito: Mitochondrial targeting sequence from the subunit VIII of human cytochrome c oxidase, H2B: Histone 2B, MAP4: microtubule-associated protein 4, Lyn11: membrane localization signal. Scale bars 10 μm.

In terms of spectral properties, the emission peak of FAST could be tuned from 540 nm to 600 nm by changing the substituents on the aromatic ring of the HBR scaffold^13^ (**Supplementary Figure 1**). To shift the spectral properties to the far-red, we used a coordinated strategy of fluorogen engineering and protein engineering to develop frFAST, a FAST variant able to recognize HPAR-3OM. Similarly to HBR derivatives, HPAR-3OM exhibits essentially undetectable fluorescence in solution or in cells, but displays red-shifted spectral properties because of an extra double bond between the phenol and the rhodanine moieties which elongates the π-electron conjugation. We demonstrated the efficiency of frFAST for multicolor imaging in cells and chicken embryo, and for *in vivo* imaging. frFAST was furthermore used to design a far-red fluorescent reporter for the detection of rapid and transient protein-protein interactions.

## RESULTS

### Engineering and characterization of frFAST

HBR analogs are non-fluorescent in solution, but they fluoresce strongly when immobilized within FAST due to inaccessible radiationless decay channels. Stabilization of their phenolate state within FAST red-shifts their absorption. This spectral change enables one to selectively excite FAST without exciting free chromophores protonated at physiological pH, further increasing the fluorogenic effect. It is assumed that FAST stabilized the phenolate state of HBR analogs via the same network of hydrogen bonds that stabilizes the phenolate state of the hydroxycinnamoyl chromophore within the parental PYP^12,16^.

In order to push the spectral properties further to the red edge of the visible spectrum, we added a double bond between the electron-donating phenol and the electron-withdrawing rhodanine heterocycle. The 4-hydroxyphenylallylidene rhodanine (HPAR) was previously shown to display red-shifted absorption and emission because of extended π-electron conjugation^21^. We generated binders able to activate the fluorescence of HPAR using a library of 10^6^ variants of FAST displayed on yeast cells. This library was incubated with HPAR and screened by fluorescence-activating cell sorting. Iterative rounds of sorting allowed us to identify a variant with the mutations D71V and P73T that formed a fluorescent complex with HPAR, characterized by 500 nm / 635 nm absorption-emission peaks and a fluorescence quantum yield ϕ = 8 %.

We tested the properties of FAST^D71V,P73T^ with various HPAR analogs, and discovered that HPAR-3OM, which bears an additional methoxy group on the aromatic ring, formed a complex with FAST^D71V,P73T^ displaying further red-shifted absorption-emission peaks (528 nm / 663 nm) and a higher fluorescence quantum yield, ϕ = 13 %. This was consistent with our previous observations that modifications to the electron-donating phenol moiety could redshift the absorption and emission of this class of pushpull molecules^13^. We thus used directed evolution to further improve the properties of FAST^D71V,P73T^ with HPAR-3OM. A library of 6 × 10^7^ variants was prepared by random mutagenesis and displayed on yeast cells. Iterative rounds of sorting in presence of HPAR-3OM allowed us to select an improved variant, which was further refined by introduction of the mutation V107I previously described to increase FAST brightness^16^. The resulting variant, possessing the mutations F62L, D71V, P73S, E74G and V107I relative to FAST, binds HPAR-3OM tightly with a *K*_D_ of 1 μM (**Supplementary Figure 2**), and strongly activates its far-red fluorescence (**Figure 1b**), forming a fluorescent complex with 555 nm / 670 nm absorption-emission peaks, a fluorescence quantum yield ϕ = 21%, and a molar absorptivity ε = 45,000 M^−1^.cm^−1^. HPAR-3OM undergoes a 130 nm red shift in absorption upon binding, in accordance with a change of ionization state from protonated in solution to deprotonated when bound (**Figure 1b** left), ensuring that unbound HPAR-3OM is fully invisible (because it does not absorb) at the wavelength used for exciting the complex. The width of the absorption band allows efficient excitation with the 543 nm, 594 nm and 633 nm lines of helium-neon lasers or the 561 nm line of yellow diode lasers. Because of the far-red fluorescence of the complex and the red-shift in absorption undergone by HPAR-3OM upon complexation, this improved variant was designated *far-red fluorescence activating and absorption shifting tag* (frFAST). With a large Stokes shift (= 115 nm), frFAST is singular among the far-red fluorescent proteins, which usually display narrower Stokes shifts, offering thus new possibilities for biological imaging. Note that frFAST conserves the ability to bind HBR analogs, although with lower performances than FAST variants (**Supplementary Table 1**).

### Imaging frFAST in live cells

We next asked whether frFAST could be used to encode far-red fluorescence in living cells. We first verified that HPAR-3OM had no deleterious effects on cells at the concentrations used for labeling. We exposed HeLa cells to 10 μM HPAR-3OM for 24 hours. No effects on cell health and viability were observed (**Supplementary Figure 3**). We next expressed frFAST in HeLa and U2OS cells, and treated cells with 10 μM HPAR-3OM. Robust far-red fluorescence was seen in transfected cells after 15 seconds of incubation. Furthermore, expression of frFAST was homogenous and no intracellular aggregates were observed (**Figure 1c**).

We then verified that frFAST could be used as a tag for labeling intracellular proteins. We expressed frFAST protein fusions with the histone H2B (H2B–frFAST), the mitochondrial targeting sequence from the subunit VIII of human cytochrome c oxidase (Mito–frFAST), and the microtubule-associated protein (MAP) 4 (MAP4–frFAST) in HeLa and U2OS cells. Treatment with HPAR-3OM led to far-red fluorescence in the expected localizations, demonstrating that frFAST fusions were functional and exhibited correct behavior (**Figure 1c**).

We next tested the photostability of frFAST:HPAR-3OM. Long-term imaging of HeLa cells expressing frFAST fused to H2B allowed us to show that frFAST displayed high photostability (**Supplementary Figure 4**). In direct comparison with smURFP^5^ and iRFP670^3^ – two dimeric far-red fluorescent proteins having emission properties comparable to frFAST – frFAST appeared a bit less photostable than iRFP670 but more than smURFP (**Supplementary Figure 4**), which was previously reported to be more photostable than most monomeric far-red fluorescent proteins^6^.

We next evaluated the use of frFAST for multi-color imaging. We imaged live HPAR-3OM-treated mammalian cells co-expressing frFAST, EGFP and mCherry fusions using a confocal microscope with adjustable emission bandwidth (**Figure 1d**). All fusions showed the expected localizations and clear spectral separation of their fluorescence signals, which allowed us to perform three-color imaging in live cells. This set of experiments showed that frFAST could be used as an additional reporter for highly multiplexed imaging.

### Imaging protein-protein interactions in live cells

One of the advantages of the FAST system is that its reversible interaction with HBR analogs permitted the development of splitFAST, a split fluorescent reporter with reversible complementation for the visualization of dynamic protein-protein interactions^20^. We thus examined whether this property was retained by frFAST, despite the changes to the protein that allow it to recognize this class of larger fluorogen. Two complementary fragments were generated by splitting frFAST in its last loop between Ser 114 and Gly 115 (**Figure 2a**). To evaluate the ability of the two fragments to complement when fused to two interacting proteins, C-frFAST (fragment 115-125) and N-frFAST (fragment 1-114) were fused respectively to the FK506-binding protein (FKBP) and the FKBP-rapamycin-binding domain of mammalian target of rapamycin (FRB), two proteins that interact together in the presence of rapamycin. HEK 293T cells co-expressing FRB-N-frFAST and FKBP-C-frFAST were treated with 10 μM HPAR-3OM. Induction of FRB-FKBP dimerization by addition of rapamycin led to an increase of the far-red fluorescence, in accordance with an interaction-dependent complementation of the two split fragments (**Figure 2b-d**). The median fluorescence fold increase was 7-fold (**Figure 2c**). Time-lapse imaging showed that complementation occurred within a few minutes after addition of rapamycin (**Figure 2b,d**), in agreement with the rapid formation of the FRB-FKBP-rapamycin complex.

**Figure 2.**
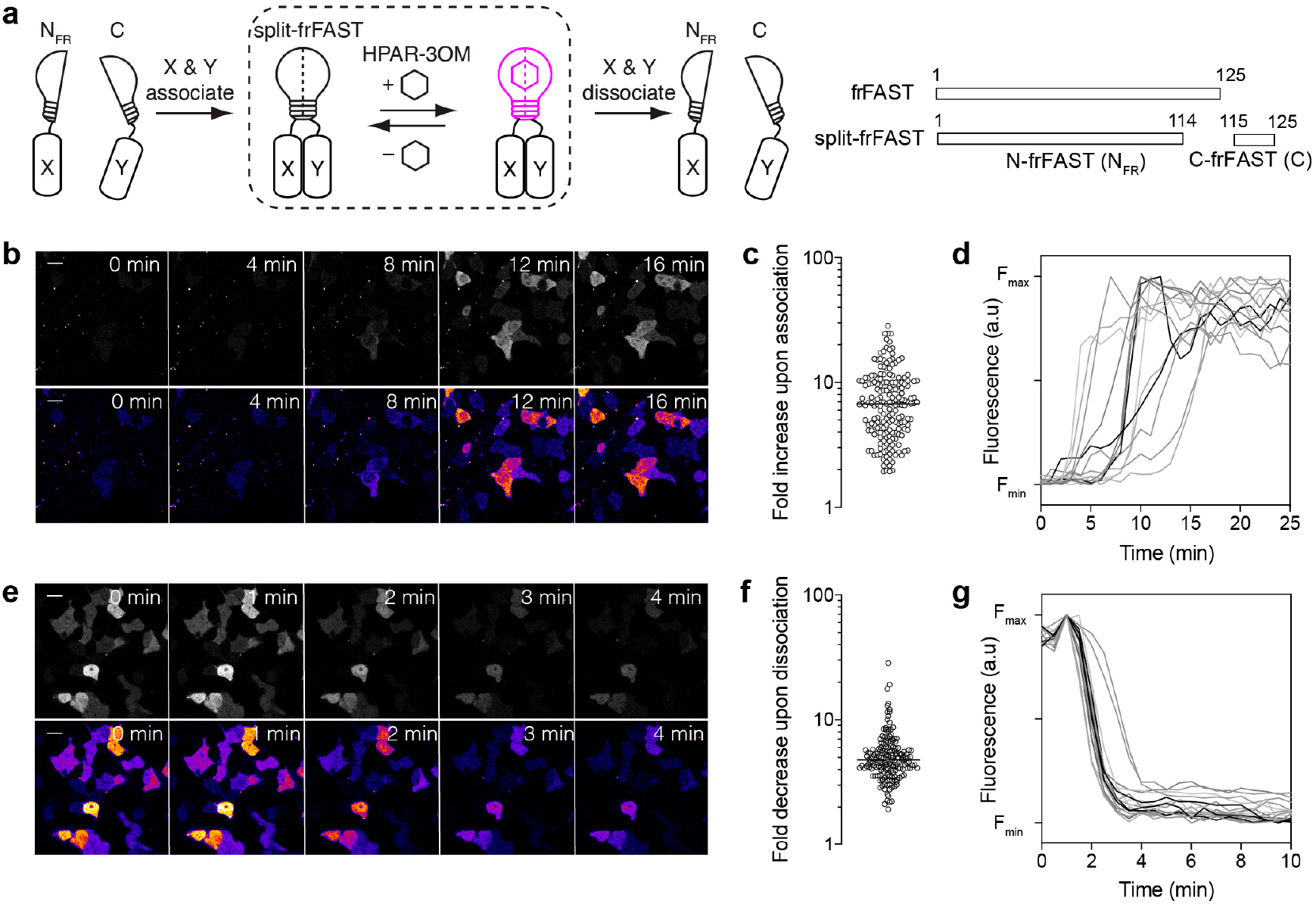
Design of a far-red fluorescent reporter for the detection of dynamic protein-protein interactions. **(a)** Principle and design of split-frFAST. **(b-d)** HEK293T cells co-expressing FK506-binding protein (FKBP)-C-frFAST and FKBP-rapamycin-binding domain of mammalian target of rapamycin (FRB)-N-frFAST were labeled with 10 μM HPAR-3OM, and imaged before and after addition of 500 nM rapamycin. **(b)** Representative time-lapse. Scale bar 20 μm. **(c)** Fluorescence fold increase upon FKBP-FRB association, n = 185 cells from three experiments. The line indicates the median. **(d)** Temporal evolution of the fluorescence intensity after rapamycin addition in HPAR-3OM-treated cells co-expressing FRB-N-frFAST and FKBP-C-frFAST (n = 14 cells). **(e-g)** HEK293T cells co-expressing FKBP-N-frFAST and FKBP-C-frFAST were treated with 100 nM AP1510 and labeled with 10 μM HPAR-3OM. Cells were then imaged before and after the addition of 1.1 μM rapamycin. **(e)** Representative timelapse. Scale bar 20 μm. **(f)** Fluorescence fold decrease upon FKBP-FKBP dissociation, n = 170 cells, from three experiments. The line indicates the median. **(g)** Temporal evolution of the fluorescence intensity after rapamycin addition in AP1510-treated cells co-expressing FKBP-N-frFAST and FKBP-C-frFAST (n = 18 cells).

We next tested the reversibility of split-frFAST using the ability of rapamycin to dissociate an AP1510-induced FKBP homodimer. We co-expressed FKBP-N-frFAST and FKBP-C-frFAST in HEK 293T cells. Cells were pre-treated with AP1510 for 2 h to form the FKBP homodimer and 10 μM HPAR-3OM was added to visualize the complemented split-frFAST. The addition of rapamycin led to a median fluorescence fold decrease of 5-fold, in agreement with a disassembly of split-frFAST concomitant with the dissociation of the FKBP homodimer (**Figure 2e-g**). Time-lapse imaging showed that complementation was reversed within a few minutes (**Figure 2e,g**), in agreement with the rapid disassembly of FKBP homodimer.

Overall, this set of experiments suggested that split-frFAST could be used to monitor protein complex formation and dissociation with high spatial and temporal resolution.

### Imaging frFAST in live zebrafish embryos and larvae

The use of far-red fluorescent tags is particularly important when working with multicellular organisms, due to the lower autofluorescence and the better light penetration in tissues at longer wavelengths. We thus examined the performance of frFAST in multicellular organisms. As model, we first used zebrafish embryo and larvae. We initially tested the toxicity of HPAR-3OM and frFAST. Zebrafish embryos were incubated with 5 μM HPAR-3OM during 1 hour at 50% epiboly, or overnight from 50% epiboly to 24 hpf. In both cases, no significant deleterious effects were observed at 48 hpf (**Supplementary Figure 5a**). The effect of expressing an frFAST fusion was tested by injecting mRNA coding for H2B-frFAST in zebrafish embryos at the one-cell stage. Embryos developed correctly and no deleterious effects were observed at 24 hpf (**Supplementary Figure 5b**). As a comparison, we observed that the expression of H2B fused to smURFP was highly toxic, and led to 50% dead embryos and 50% embryos with strong axis defects at 24 hpf.

We then evaluated the labeling efficiency in zebrafish embryos. We injected mRNA coding for H2B-frFAST in zebrafish embryo at the one-cell stage, together with an mRNA encoding H2B-EGFP to account for injection efficiency. Embryos were imaged after treatment with 5 μM HPAR-3OM for 30 min. We observed that expression of H2B-frFAST was efficiently detected during gastrulation (**Figure 3a,b**) and at 24 hpf (**Figure 3c,d**). Similar experiments using H2B fused to the far-red fluorescent protein iRFP713^2^ revealed that iRFP713 could only be detected at 24 hpf but not during gastrulation (**Figure 3e-h**). The absence of iRFP713 signal during gastrulation might be due to the slow incorporation of biliverdin and/or the lack of biliverdin at this development stage. Overall, these results demonstrated that frFAST could be an efficient far-red fluorescent tag for imaging proteins in zebrafish at various stages of embryogenesis, even early ones.

**Figure 3.**
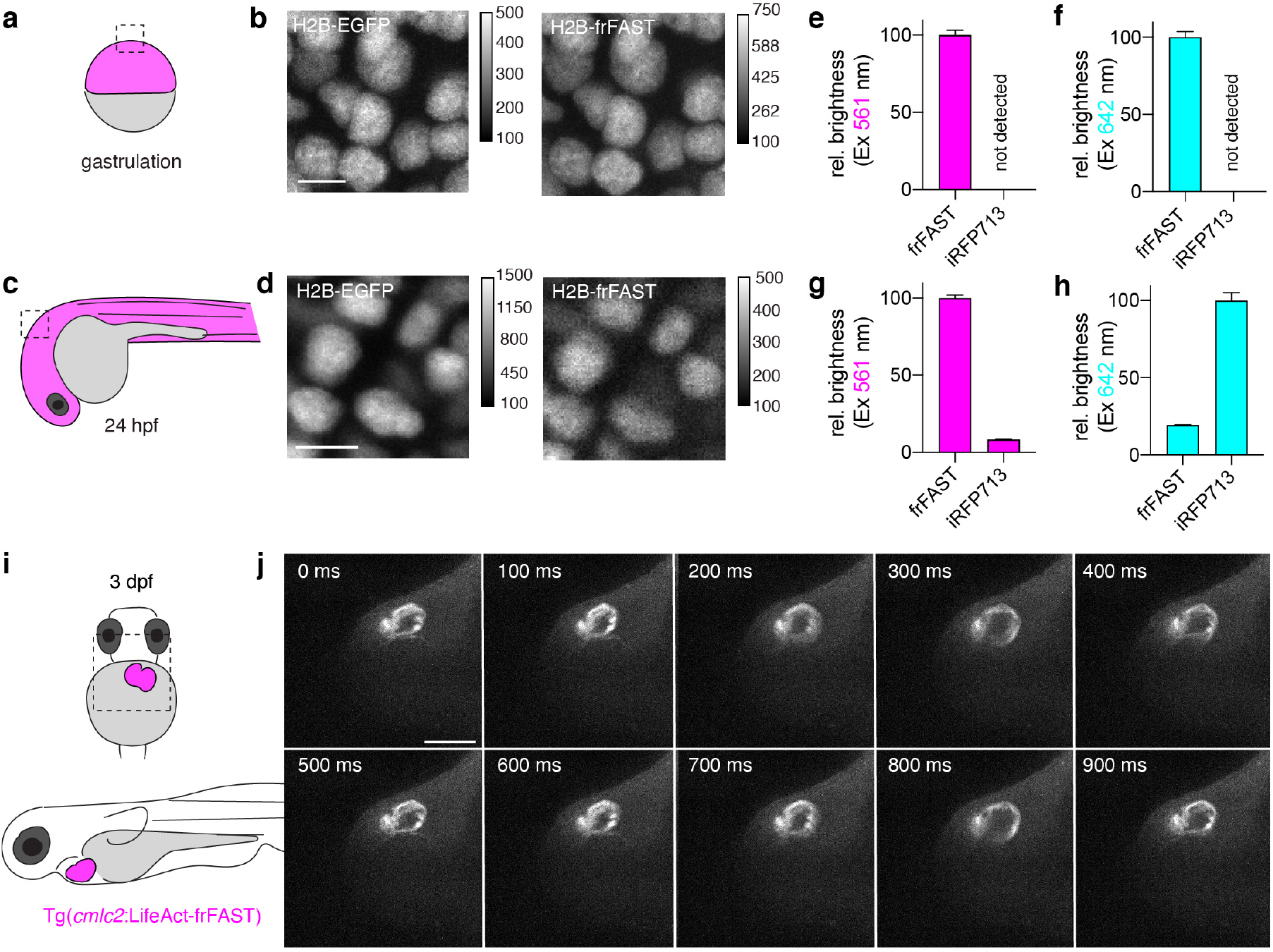
frFAST enables to encode far-red fluorescence in zebrafish embryos and larvae. **(a-d)** mRNA encoding H2B-EGFP and H2B-frFAST were micro-injected in zebrafish embryo at the one-cell stage. The confocal micrographs shows few cells in zebrafish embryos during gastrulation **(a,b)** and at 24 hpf **(c,d)**. Embryos were incubated with 5 μM HPAR-3OM for 30 min prior imaging. Micrographs were recorded with a spinning-disk confocal microscope with the following settings: EGFP Ex 491 nm / Em 497-547 nm, frFAST Ex 561 nm / Em LP 655 nm. Scale bars 10 μm. **(e-h)** Effective brightness of frFAST and iRFP713 during gastrulation **(e,f)** and at 24 hpf **(g,h)** exciting at 561 nm (e,g) or 642 nm **(f,h)**. frFAST and iRFP713 were expressed in zebrafish embryo as fusion to H2B, and fluorescence was quantified by confocal microscopy. The relative brightness were determined after normalization to fluorescence of co-expressed H2B-EGFP. Brightness represents the mean ± sem determined from n > 50 cells. **(i,j)** Time-lapse imaging of a 3 dpf transgenic zebrafish larva expressing LifeAct-frFAST in the myocardium (see also **Supplementary Movie 1**). The larva was incubated with 5 μM HPAR-3OM for 2 h prior imaging. Micrographs were recorded with a spinning-disk confocal microscope with the following settings: frFAST Ex 561 nm / Em LP 655 nm. Scale bar 100 μm.

We next tested if the far-red fluorescence of frFAST could be detected in internal organs in zebrafish larvae. We established a transgenic line of zebrafish with expression of frFAST fused to LifeAct, a peptide binding filamentous actin, under the control of the *cmlc2* promoter, known to be sufficient to drive myocardium-specific expression (**Figure 3i**). Labeling with HPAR-3OM and imaging with a high speed-spinning disk confocal microscope allowed us to observe the myocardium in beating heart in 3 dpf larvae (**Figure 3j** and **Supplementary Movie 1**). This experiment showed that frFAST fusion could be efficiently labeled and detected in internal organs in live zebrafish larvae.

### Imaging frFAST in chicken embryos

To verify that frFAST could be used in other multicellular organisms, we used chicken embryo as a second model. Plasmids encoding frFAST or miRFP670nano^6^ (fused to H2B) under the control of the CAGGS promoter^22^ were each electroporated in one of each side of the neural tube *in ovo*, at embryonic day 2 (E2, HH stage 13-14) (**Figure 4a,b**). An EGFP reporter (fused to a membrane localization signal) was co-injected with each construct to monitor electroporation efficiency. 24h later, embryos with homogeneous bilateral EGFP expression in the neural tube were dissected and imaged in absence and in presence of HPAR-3OM (**Figure 4c**). Only cells expressing frFAST lighted up upon addition of HPAR-3OM, demonstrating the selective labeling of frFAST in chicken embryos. In addition, cells located in the side electroporated with the plasmid encoding frFAST showed higher fluorescence than cells located in the side electroporated with the plasmid encoding miRFP670nano when excited at 640 nm, suggesting that frFAST outperformed miRFP670nano in terms of effective brightness in this context. Time-lapse imaging furthermore showed that maximal fluorescence was achieved 20-25 min after HPAR-3OM addition (**Supplementary Movie 2**), demonstrating that HPAR-3OM efficiently diffused across the embryo.

**Figure 4.**
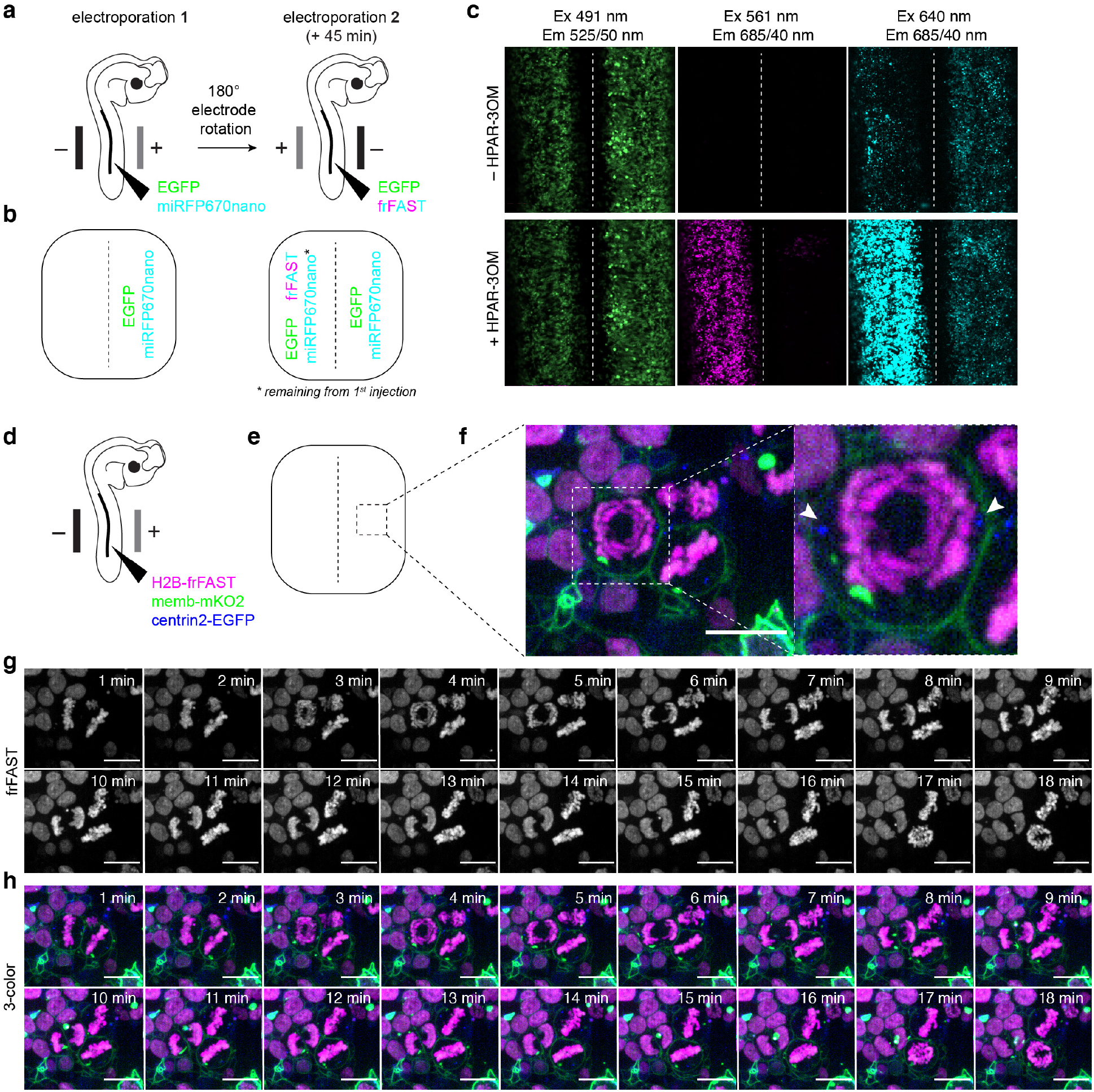
frFAST enables to encode far-red fluorescence in chicken embryos. **(a-c)** Plasmids encoding H2B-miRFP670nano and H2B-frFAST were each electroporated in one of each side of the neural tube in ovo at embryonic day 2 (E2, HH stage 13-14) **(a)**. An EGFP reporter (fused to a membrane localization signal) was co-injected with each construct to monitor electroporation efficiency. 24 h later, embryos with homogeneous bilateral EGFP expression in the neural tube were dissected **(b)**, and imaged in absence and in presence of 10 μM HPAR-3OM using a spinning-disk confocal microscope **(c)** (see **Supplementary Movie 2** for time-lapse imaging of the labeling). Note that before addition of HPAR-3OM, some miRFP670nano signal is detected on the left side of the embryo. This is caused by residual plasmid from the first injection remaining in the neural tube at the time of the second injection. **(d-h)** Multicolor imaging. Plasmids encoding H2B-frFAST, mKO2 (fused to a membrane localization signal) and Centrin2-EGFP were electroporated in the neural tube in ovo, at embryonic day 2 (E2, HH stage 13-14) **(d,e)**. 24 h later, embryos were dissected, and imaged in presence of 10 μM HPAR-3OM using a spinning-disk confocal microscope. **(f)** Three-color micrographs showing the correct localization of the three proteins (Centrin2-EGFP is indicated with white arrows). **(g,h)** Time-lapse showing cell division. Micrographs were recorded with the following settings: EGFP Ex 491 nm / Em 525/50 nm, mKO2 Ex 561 nm / Em 617/73 nm, frFAST Ex 561 nm / Em 685/40 nm. Scale bar 10 μm.

We next tested our ability to perform multicolor imaging in the chicken embryo. We electroporated plasmids encoding frFAST (fused to H2B) together with EGFP (fused to centrin2) and mKO2 (fused to a membrane localization signal) in the neural tube as previously (**Figure 4d**). En-face time-lapse imaging of the neuroepithelium with a spinning-disk confocal microscope allowed us to visualize the three proteins (**Figure 4f**) and follow their dynamics during cell division (**Figure 4g,h**). This set of experiments allowed us to further validate the use of frFAST for multicolor imaging of dynamic processes with a subcellular resolution in multicellular organisms.

## DISCUSSION

A coordinated strategy of protein engineering and fluorogen engineering allowed us to generate a small far-red fluorescent chemogenetic reporter named frFAST, which forms a bright far-red fluorescent complex with the fluorogenic chromophore HPAR-3OM. HPAR-3OM is essentially non-fluorescent in solution and in cells, allowing one to image proteins fused to frFAST with high contrast in the presence of free fluorogenic ligand. The high fluorogenicity and exquisite labeling specificity allowed the imaging of proteins not only in mammalian cells but also in multicellular systems such as zebrafish embryo/larvae and chicken embryo. With 555 nm / 670 nm absorption-emission peaks, frFAST is a far-red fluorescent reporter with a large Stokes shift, making it singular in the landscape of far-red fluorescent proteins, which usually display narrow Stokes shifts (**Supplementary Table 2**), expanding thus the fluorescence toolkit with new properties and possibilities. Its far-red emission enabled us to perform multicolor imaging both in cells and in multicellular systems.

In contrast with biliverdin-based far-red fluorescent proteins, which display slow fluorescence maturation (half-time > 100 min) because of the need to covalently bind biliverdin, frFAST binds HPAR-3OM almost instantaneously. Combined with the high cellular uptake of HPAR-3OM, this allows the labeling to occur with only few tens of seconds in mammalian cells and few tens of minutes in zebrafish embryos/larvae and chicken embryos. As frFAST requires the addition of an exogenous fluorogen, its brightness can be maximized by adjusting the HPAR-3OM concentration to have almost 100% fluorescent complex, avoiding the problems encountered with far-red fluorescent proteins, in which effective brightness can be limited by the availability and accessibility of biliverdin^7^. frFAST could for instance be visualized during gastrulation, a development stage where iRFP713 was not detectable in zebrafish. The small size of frFAST is another advantage. Small monomeric tags are desirable as they interfere less with the labeled proteins, and have minimal genetic footprint. The toxicity we observed in zebrafish when using the far-red fluorescent protein smURFP illustrates how fluorescent proteins can affect the function of labeled proteins. With a molecular weight of less than 14 kDa, frFAST is smaller than the smallest monomeric far-red fluorescent protein, miRFP670nano^6^, a 17 kDa protein engineered from a cyanobacteriochrome. Finally, with a fluorescence quantum yield of 21% and a molar absorptivity of 45,000 M^−1^cm^−1^, the molecular brightness (i.e. the product of the fluorescence quantum and the molar absorptivity) of frFAST equals or exceeds that of most monomeric far-red fluorescent proteins (**Supplementary Table S2**).

As a very close structural and functional relative of the chemogenetic reporter FAST, frFAST displays the same advantages. The need to add HPAR-3OM to generate the far-red fluorescence allows the design of experimental protocols requiring on-demand labeling. In addition, as no molecular oxygen is required for generating the far-red fluorescence, frFAST should enable the detection of fusion proteins in anaerobic conditions. Finally, as FAST, frFAST should be well suited as optical reporting module for the design of various biosensors. We demonstrated in this work that frFAST could be turned, using the design previously used for the generation of splitFAST^20^, into a far-red split fluorescent reporter with rapid and reversible complementation for the detection of protein-protein interactions with no additional engineering. We anticipate that frFAST could be used for the design of far-red fluorescent biosensors through allosteric coupling of frFAST (or circularly permuted versions) and analyte binding domains, in the same way FAST was previously used for the design of biosensors^19^.

## NOTES

The authors declare the following competing financial interest: A.G. is co-founder and hold equity in Twinkle Bioscience / The Twinkle Factory, a company commercializing the FAST technology.

## ACKNOWLEDGEMENT

We thank K. D. Wittrup, for providing us with the pCTCON2 vector and the EBY100 yeast strain for the yeast display selection. We also thank the flow cytometry facility CISA (Cytométrie Imagerie Saint-Antoine) of UMS LUMIC at the Faculty of Medicine of Sorbonne Université, and, more particularly, Annie Munier for her assistance. This work has been supported by the European Research Council (ERC-2016-CoG-724705 FLUOSWITCH).

## SUPPORTING INFORMATION

Supporting information contains the **Supplementary Movies 1-2**, the **Supplementary Tables 1-2** and the **Supplementary Figures 1-5**.

## MATERIALS AND METHODS

### General

Synthetic oligonucleotides used for cloning were purchased from Sigma Aldrich or Integrated DNA Technology. PCR reactions were performed with Q5 polymerase (New England Biolabs) in the buffer provided. PCR products were purified using QIAquick PCR purification kit (Qiagen). The products of restriction enzyme digests were purified by preparative gel electrophoresis followed by QIAquick gel extraction kit (Qiagen). Restriction endonucleases, T4 ligase, Phusion polymerase, Taq ligase, and Taq exonuclease were purchased from New England Biolabs and used with accompanying buffers and according to manufacturer protocols. Isothermal assemblies (Gibson assembly) were performed using homemade mix prepared according to previously described protocols^23^. Small-scale isolation of plasmid DNA was done using QIAprep miniprep kit (Qiagen) from 2 mL of overnight culture. Large-scale isolation of plasmid DNA was done using the QIAprep maxiprep kit (Qiagen) from 150 mL of overnight culture. All plasmid sequences were confirmed by Sanger sequencing with appropriate sequencing primers (GATC services - Eurofins Genomics).

### Sequences

Sequence of frFAST (125 amino acids, MW = 13,788.7 Da)

MEHVAFGSEDIENTLAKMDDGQLDGLAFGAIQLDGDGNILQYNAAEGDITGRDPKQVIGKNLFKDVAPGTVSSGFYGKFKEGVASGNLNTMFEWMIPTSRGPTKVKIHMKKALSGDSYWVFVKRV

DNA sequence coding for frFAST (375 bp)

atggagcatgttgcctttggcagtgaggacatcgagaacactttggccaaaatggacgacggacaactggatgggttggcctttggcgcaattcagctcgatggtgacgggaatatcctgcagtacaatgctgctgaaggagacatcacaggcagagatcccaaacaggtgattgggaagaacttattcaaggatgttgcacctggaacggtttcctccgggttttacggcaaattcaaggaaggcgtagcgtcagggaatctgaacaccatgttcgaatggatgataccgacaagcaggggaccaaccaaggtcaagatacacatgaagaaagccctttccggtgacagctattgggtctttgtgaaacgggtg

Sequence of N-frFAST (114 amino acids, MW = 12,251.15 Da)

MEHVAFGSEDIENTLAKMDDGQLDGLAFGAIQLDGDGNILQYNAAEGDITGRDPKQVIGKNLFKDVAPGTVSSGFYGKFKEGVASGNLNTMFEWMIPTSRGPTKVKIHMKKALS

DNA sequence coding for N-frFAST (342 bp)

atggagcatgttgcctttggcagtgaggacatcgagaacactttggccaaaatggacgacggacaactggatgggttggcctttggcgcaattcagctcgatggtgacgggaatatcctgcagtacaatgctgctgaaggagacatcacaggcagagatcccaaacaggtgattgggaagaacttattcaaggatgttgcacctggaacggtttcctccgggttttacggcaaattcaaggaaggcgtagcgtcagggaatctgaacaccatgttcgaatggatgataccgacaagcaggggaccaaccaaggtcaagatacacatgaagaaagccctttcc

Protein sequence of C-frFAST (11 amino acids, MW 1,355.6 Da)

GDSYWVFVKRV

DNA sequence coding for C-frFAST (33 bp)

ggtgacagctattgggtctttgtgaaacgggtg

### Fluorogens

The synthesis of HPAR-3OM was performed using commercially available reagents used as received. To a stirred solution of rhodanine (133 mg, 1.0 mmol) and 4-Hydroxy-3-methoxycinnamaldehyde (178 mg, 1.0 mmol) in ethanol (10 mL) was added 4- (dimethylamino)pyridine (12 mg, 0.1 mmol). The solution was stirred at reflux for 20 h. Acetic acid was added to neutralize the solution. After cooling at 4°C overnight, the precipitate was filtered. The crude solid was washed with diethyl ether and dried. HPAR-3OM was obtained pure as a red powder (150 mg, 52% yield). ^1^H NMR (300 MHz, DMSO-d6, δ in ppm) 13.54 (s, 1H), 9.69 (s, 1H), 7.33 (m, 2H), 7.23 (d, *J* = 15 Hz, 1H), 7.08 (d, *J* = 8.1 Hz, 1H), 6.84 (m, 2H), 3.84 (s, 3H). ^13^C NMR (75 MHz, DMSO-d6, δ in ppm) 194.7, 168.6, 151.6, 146.6, 133.9, 130.0(2C), 123.1, 122.3, 118.3, 111.9(2C), 39.7. HRMS (ESI): m/z 294.0253 [M+H]^+^, calcd mass for [C_13_H_12_NO_3_S_2_]^+^: 294.0259. ^1^H and ^13^C NMR spectra were recorded at room temperature on a Bruker AM 300 spectrometer. Mass spectrometry analysis was performed by the Institut de Chimie Organique et Analytique de l’Université d’Orléans.

The preparation of HPAR (4-hydroxy-phenylallylidene rhodanine) was previously described^21^. The preparation of HMBR, HBR-3,5DM, HBR-3OM, HBR-3,5DOM was previously described^12,13^. HMBR, HBR-3,5DM, HBR-3OM, HBR-3,5DOM are commercially available from The Twinkle Factory (the-twinkle-factory.com) under the name ^TF^Lime, ^TF^Amber, ^TF^Citrus and ^TF^Coral, respectively.

### Directed evolution

#### Library construction

The random mutagenesis library of FAST variants used for the initial yeast display screen with HPAR was previously described^24^. For the library of FAST^D71V,P73T^ variants, the gene of FAST^D71V,P73T^ was randomly mutagenized by error-prone PCR using the GeneMorph II Random Mutagenesis Kit (Agilent). The PCR product was digested with NheI and BamHI, and then ligated into the pCTCON2 vector using NheI / BamHI restriction sites. Large-scale transformation into DH10B *E. coli* cells was performed by electroporation. The DNA was then purified and retransformed into EBY100 yeast strain using a large-scale high-efficiency transformation protocol^25^. The final library contains 6 × 10^7^ clones.

#### FACS selection

The yeast library (about 1 × 10^9^ cells) was grown overnight (30°C, 280 rpm) in 1 L of SD (20 g/L dextrose, 6.7 g/L yeast nitrogen base, 1.92 g/L yeast synthetic dropout without tryptophane, 7.44 g/L NaH_2_PO_4_ and 10.2 g/L Na_2_HPO_4_-7H2O, 1% penicillin-streptomycin 10,000 U/mL). Yeast culture was then diluted to OD_600_ 1 in 1L of SD and grown (30°C, 280 rpm) until OD_600_ 2-5. Next, 5 × 10^9^ cells yeast cells were then collected and grown for 36 h (23°C, 280 rpm) in 1L SG (20 g/L galactose, 2 g/L dextrose, 6.7 g/L yeast nitrogen base, 1.92 g/L yeast synthetic dropout without tryptophane, 7.44 g/L NaH_2_PO_4_ and 10.2 g/L Na_2_HPO_4_-7H2O, 1% penicill-instreptomycin 10,000 U/mL). 5 × 10^8^ induced cells were then pelleted by centrifugation (25°C, 3 min, 2,500 g), washed with 10 mL DPBS-BSA (137 mM NaCl, 2.7 mM KCl, 4.3 mM Na_2_HPO_4_, 1.4 mM KH2PO4, 1 g/L bovine serum albumin, pH 7.4). Cells were incubated for 30 min at room temperature in 200 μL of 1/250 primary antibody chicken anti-Myc IgY (Life Technologies) solution in DPBS-BSA. Cells were then washed with 10 mL DPBS-BSA, and incubated in 200 μL of 1/100 secondary antibody Alexa Fluor^®^ 647–goat anti-chicken IgG (Life Technologies) solution in DPBS-BSA for 20 min on ice. Cells were then washed with DPBS. For the selection of HPAR binders using the library of FAST variants, cells were incubated in 10 mL DPBS supplemented with 10 μM HPAR, and sorted on a MoFlo™ Astrios Cell Sorter (Beckman Coulter) equipped with a 488 nm and a 640 nm laser. The fluorescence of HPAR binders was detected using the following parameters: Ex 488 nm, Em 620 ± 29 nm. For the selection of HPAR-3OM binders using the library of FAST^D71V,P73T^ variants, cells were incubated in 10 mL DPBS supplemented with 5 μM HPAR-3OM, and sorted on a MoFlo™ Astrios Cell Sorter (Beckman Coulter) equipped with a 561 nm laser and a 640 nm laser. The fluorescence of HPAR-3OM binders was detected using the following parameters: Ex 561 nm, Em 664 ± 11 nm. For both selections, the sorted cells were collected in SD, grown overnight (30°C, 240 rpm) and spread on SD plates (SD supplemented with 182 g/L sorbitol, 15 g/L agar). Plates were incubated for 60 h at 30°C. The cell lawn was collected in SD supplemented with 30% glycerol, aliquoted and frozen or directly used in the next round. After FACS selection, clones were screened by flow cytometry and their DNA was isolated using a miniprep kit (Qiagen), transformed into DH10B and re-isolated for sequencing.

### Characterization of variants

#### Cloning

Plasmids driving bacterial expression of the variants with an N-terminal His-tag under the control of a T7 promoter were constructed by inserting the encoding sequence between NheI and XhoI restriction sites in the pET28a vector.

#### Expression

The plasmids were transformed in Rosetta(DE3)pLysS E. coli (New England Biolabs). Cells were grown at 37°C in Lysogeny Broth (LB) medium complemented with 50 μg/ml kanamycin and 34 μg/ml chloramphenicol to OD_600nm_ 0.6. Expression was induced at 37°C for 4 h by adding isopropyl β-D-1-thiogalactopyranoside (IPTG) to a final concentration of 1 mM. Cells were harvested by centrifugation (4,300 × g for 20 min at 4°C) and frozen.

#### Purification protocol 1

The cell pellet was resuspended in lysis buffer (phosphate buffer 50 mM, NaCl 150 mM, MgCl2 2.5 mM, protease inhibitor, DNase, pH 7.4) and sonicated (5 min at 20 % of amplitude). The lysate was incubated for 2 h at 4 °C to allow DNA digestion by DNase. Cellular fragments were removed by centrifugation (9,300 × g for 1h at 4°C). The supernatant was incubated overnight at 4°C under gentle agitation with Ni-NTA agarose beads in phosphate buffered saline (PBS) (sodium phosphate 50 mM, NaCl 150 mM, pH 7.4) complemented with 10 mM Imidazole. Beads were washed with 20 volumes of PBS containing 20 mM Imidazole, and with 5 volumes of PBS complemented with 40 mM Imidazole. His-tagged proteins were eluted with 5 volumes of PBS complemented with 0.5 M Imidazole, followed by dialysis with PBS.

#### Purification protocol 2

The cell pellet was resuspended in 1× Tris-EDTA-sucrose (TES) buffer and incubated for 1 hr. The lysate was then diluted by three using 0.25 × TES buffer and incubated for 45 min. Cellular fragments were removed by centrifugation (9200 × g for 1.5 h at 4°C). The supernatant was incubated overnight at 4°C under gentle agitation with Ni-NTA agarose beads in phosphate buffered saline (PBS) (sodium phosphate 50 mM, NaCl 150 mM, pH 7.4) complemented with 10 mM imidazole. Beads were washed with ~20 volumes of PBS containing 20 mM imidazole, and with ~5 volumes of PBS complemented with 40 mM imidazole. His-tagged proteins were eluted with ~5 volumes of PBS complemented with 0.5 M imidazole. The buffer was exchanged to PBS (50 mM phosphate, 150 mM NaCl, pH 7.4) using PD-10 desalting columns. In both protocols, purity of the proteins was evaluated using SDS-PAGE electrophoresis stained with Coomassie blue.

#### Spectroscopic characterization

Steady state UV-Vis absorption spectra were recorded using a Cary 300 UV-Vis spectrometer (Agilent Technologies), equipped with a Versa20 Peltier-based temperature-controlled cuvette chamber (Quantum Northwest) and fluorescence data were recorded using a LPS 220 spectrofluorometer (PTI, Monmouth Junction, NJ), equipped with a TLC50TM Legacy/PTI Peltier-based temperature-controlled cuvette chamber (Quantum Northwest). Fluorescence quantum yield were determined as previously described using FAST:HBR-3,5DOM as reference^16^.

#### Binding affinity

Thermodynamic dissociation constants were determined as previously described^12^ using a Spark 10M plate reader (Tecan) and Prism 6 to fit the data with a one-site specific binding model.

### Molecular biology

The plasmids pAG153 encoding FKBP-CFAST11 (= FKBP-C-frFAST) was previously described^20^. The plasmids pAG29, pAG28, CV2326 encoding EGFP, Lyn11-EGFP, Mito-mCherry were previously described^13,26^. The plasmids pAG504, pAG505, pAG506 allowing the mammalian expression of frFAST, H2B-frFAST, mito-frFAST were obtained by replacing the sequence coding for FAST in the previously described^12^ pAG104, pAG109, pAG156 with that coding for frFAST using standard molecular biology techniques. The plasmid pAG324 allowing the mammalian expression of H2B-mCherry was obtained by replacing the sequence coding for dronpa2 in the previously described pAG123 with that coding for mCherry. The plasmid pAG502 allowing the mammalian expression of MAP4-frFAST was obtained by replacing the sequence coding for H2B in pAG505 with that coding for MAP4 using standard molecular biology techniques. The plasmids pAG499 and pAG500 for the mammalian expression of FRB-N-frFAST and FKBP-N-frFAST were obtained by replacing the sequence coding for N-FAST in the previously described^20^ pAG148 and pAG149 with that coding for N-frFAST using Gibson assembly.

For expression in zebrafish, the plasmids #1115, #1117, #1120 and #1121 encoding H2B fusion to frFAST, EGFP, iRFP713 and smURFP respectively were obtained by substituting mCherry downstream H2B in the pT2iC6 vector (a derivative of pCS2) previously described^27^. mRNA were synthesized with a mMessage mMachine™ SP6 transcription kit (Thermofisher) according to manufacturer instructions. Plasmid #1129 used to obtain transgenic fish line expressing LifeAct-frFAST fusion under the control of the cmlc2 enhancer^28^ was constructed using the NEBuilder assembly kit (New England BioLabs) in the pT22i vector for transgenesis previously used^29^.

For expression of H2B-frFAST in the chick neural tube, the CMV promoter in pAG505 was converted to a CAGGS promoter^22^ by replacing the NdeI/BglII fragment with a NdeI/BglII CAGGS fragment from pCAGGS-MCS2 (X. Morin, unpublished). pCAGGS-H2B-miRFP670nano was obtained by converting the CMV promoter in pH2B-miRFP670nano (https://www.addgene.org/127438/) to a CAGGS promoter by replacing the NdeI/KpnI fragment with a NdeI/KpnI fragment from pCAGGS-MCS2. pCAGGS-mKO2-Farn was a kind gift of Fumio Matsuzaki.

Construction details, complete sequences and plasmids are available upon request.

### Experiments in mammalian cells

#### General

HeLa cells were cultured in Modified Eagle Medium (MEM) supplemented with 0.9 % MEM non-essential amino acids, 0.9% sodium pyruvate, and 9% (vol/vol) fetal calf serum (FCS), at 37 °C in a 5% CO_2_ atmosphere. U2OS cells were cultured in McCoy’s 5A medium supplemented with phenol red and 10% (vol/vol) FCS. HEK293T cells were cultured in Dulbecco’s Modified Eagle Medium (DMEM) supplemented with phenol red, Glutamax I, and 10% (vol/vol) FCS. For imaging, cells were seeded in μDish IBIDI (Biovalley) coated with poly-L-lysine. Cells were transiently transfected using Genejuice (Merck) according to the manufacturer’s protocol for 24 hours prior to imaging. The cells were washed with PBS, and treated with DMEM media (without serum and phenol red) containing the fluorogens at the indicated concentration. The cells were imaged directly without washing.

#### Cell viability assay

HeLa cells were treated with DMEM media containing HPAR-3OM at the indicated concentrations for the indicated times. The cell viability was evaluated by fluorescence microscopy using the LIVE/DEAD^®^ viability/cytotoxicity assay kit (Molecular Probes, Life Technologies) following the manufacturer’s protocol.

#### Sensing of protein-protein interactions

To monitor the association of the FRB-FKBP homodimer, rapamycin was added to HEK 293T cells co-expressing FRB-N-frFAST and FKBP-C-frFAST to a final concentration of 500 nM. To measure the dissociation of the FKBP-FKBP homodimer, HEK293T cells co-expressing FKBP-N-frFAST and FKBP-C-frFAST were first pre-incubated with 100 nM AP1510 for 2 hours then rapamycin was added to a final concentration of 1.1 μM for inducing the dissociation. Rapamycin was purchased from Sigma Aldrich and dissolved in DMSO to a concentration of 3 mM. AP1510 was purchased from Clontech and dissolved in ethanol to a concentration of 0.5 mM.

#### Microscopy

The confocal micrographs of mammalian cells were acquired on a Zeiss LSM 710 Laser Scanning Microscope equipped with a Plan Apochromat 63× /1.4 NA oil immersion objective. ZEN software was used to collect the data. The images were analyzed with Fiji (Image J).

### Experiments in zebrafish embryos and larvae

#### Evaluation of fluorogen toxicity

Embryos were incubated in 5 μM HPAR-3OM during 1 h at 50% epiboly or overnight from 50% epiboly to 24 hpf. Embryos with no defect, axis defects, or dead were scored at 48 hpf by brightfield microscopy.

#### Evaluation of expression toxicity

mRNA coding for H2B-EGFP (20 ng/μl) and either H2B-frFAST (80 ng/μl) or H2B-smURFP (80 ng/μl) were co-injected at the one-cell stage. Embryos with no defect, axis defects, or dead were scored at 24 hpf by brightfield microscopy.

#### Brightness characterization

mRNA coding for H2B-EGFP (20 ng/μl) and either H2B-frFAST (80 ng/μl) or H2B-iRFP713 (80 ng/μl) were co-injected at the one-cell stage. frFAST expressing embryos were incubated with 5 μM HPAR-3OM for 30 min prior imaging. Fluorescence of frFAST and iRFP713 was quantified by confocal microscopy and normalized to the fluorescence of EGFP.

#### Transgenic line

Plasmid #1129 (10 ng/μL) was co-injected with mRNA coding for Tol2 transposase (20 ng/μL) and fish were screened for germline transmission.

#### Microscopy

Imaging of embryos and larvae embeded in low-melting agarose (0.8%) was performed with a CSU-W1 Yokogawa spinning disk coupled to a Zeiss Axio observer Z1 inverted microscope equipped with a sCMOS Hamamatsu camera and a 25× (Zeiss 0.8 Imm WD: 0.19 mm) oil objective. For frFAST and iRFP713 imaging, 561 nm and 642 nm lasers, and a 655 longpass filter were used. EGFP imaging was performed using a 491nm laser and a 522/50 bandpass filter.

### Experiments in chicken embryos

JA57 chicken fertilized eggs were provided by EARL Morizeau (8 rue du Moulin, 28190 Dangers, France) and incubated at 38°C in a Sanyo MIR-253 incubator. Embryos used in this study were between E2 (HH14) and E3 (HH14 + 24 h). The sex of the embryos was not determined.

***Electroporation*** in the chick neural tube was performed at embryonic day 2 (E2, HH stage 14), by applying five pulses of 50 ms at 25 V with 100 ms in between, using a square-wave electroporator (Nepa Gene, CUY21SC) and a pair of 5-mm gold-plated electrodes (BTX Genetrode model 512) separated by a 4-mm interval. The DNA solution was injected directly into the lumen of the neural tube via glass capillaries. Bilateral electroporation was achieved by switching the electrodes polarity and repeating the procedure after 45 min. All DNA constructs were used at 0.5 μg/μl each.

***En-face culture*** of the embryonic neuroepithelium was performed at E3 (24 h after electroporation). After extraction from the egg and removal of extraembryonic membranes in PBS, embryos were transferred to 37°C F12 medium and pinned down with dissection needles at the level of the hindbrain and hindlimbs in a 35mm Sylgard dissection dish. A dissection needle was used to separate the neural tube from the somites from hindbrain to caudal end on both sides of the embryo, and the roof-plate was then slit with the needle. The neural tube and notochord were then “peeled off” from the remaining tissues and equilibrated 5 min in 1% low meting point agarose/F12 medium at 38°C. The tissue was then transferred in a drop of agarose medium to a glass-bottom culture dish (MatTek, P35G-0-14-C) and excess medium was removed so that the neural tube would flatten with its apical surface facing the bottom of the dish, in an inverted open book conformation. After 30 s of polymerization on ice, an extra layer of agarose medium was added to cover the whole tissue and hardened on ice for 1 min. 2 mL of 37°C culture medium was added (F12/Penicillin Streptomycin/Sodium pyruvate) and the culture dish was transferred to the 37°C chamber of a spinning disk confocal microscope. To image frFAST, 0.5 ml of culture medium with 50 μM HPAR-3OM (1.25 μl of a 20 mM stock in 500 μl medium) was added to the dish for a final concentration of 10 μM.

***Live imaging*** was performed on an inverted microscope (Nikon Ti Eclipse) equipped with a heating enclosure (DigitalPixel, UK), a spinning disk confocal head (Yokogawa CSU-W1), and an sCMOS Camera (Orca Flash4LT, Hamamatsu) driven by MicroManager software^30^. Image stacks were obtained at 1 min intervals either with a 10× objective (CFI Plan APO LBDA, NA 0.45, Nikon; z-step = 4 μm; Figure 4c, Movie S1) or a 100× oil immersion objective (APO VC, NA 1.4, Nikon; z-step = 1 μm; Figure 4f-h).

