## Supporting Information file for "A far-red fluorescent chemogenetic reporter for in vivo molecular imaging"

### Content

Legends of Supplementary Movies 1-2

Supplementary Tables 1-2

Supplementary Figures 1-5

Supplementary References

**Legends**

**Supplementary Movie 1. Time-lapse imaging of a 3 dpf transgenic zebrafish larva expressing LifeAct-frFAST in the myocardium.** See also **Figure 3i,j** in the main text. The larva was incubated with 5 μM HPAR-3OM for 2 h prior imaging. Micrographs were recorded with a spinning-disk confocal microscope with the following settings: frFAST Ex 561 nm / Em LP 655 nm. Scale bar 100 μm.

**Supplementary Movie 2. Labeling frFAST with HPAR-3OM in the dissected neural tube of chicken embryo.** Plasmids encoding H2B-miRFP670nano (abbreviated H2B-iRFPnano on the movie label) and H2B-frFAST were each electroporated in one of each side of the neural tube in ovo at embryonic day 2 (E2, HH stage 13-14) (see also **Figure 4a-c** in the main text). An EGFP reporter fused to a membrane localization signal (abbreviated mb-GFP on the movie label) was co-injected with each construct to monitor electroporation efficiency. 24 h later, embryos with homogeneous bilateral EGFP expression in the neural tube were dissected, and imaged upon addition of 10 μM HPAR-3OM using a spinning-disk confocal microscope (imaging settings are indicated on the movie). Note that before addition of HPAR-3OM, some miRFP670nano signal is detected on the left side of the embryo. This is caused by residual plasmid from the first injection remaining in the neural tube at the time of the second injection.

**Supplementary Table 1.** Physico-chemical properties of frFAST with HPAR-3OM and various fluorogens of the HBR family in PBS pH 7.4. Data with FAST and iFAST are also reported for comparison. Abbreviations are as follows: $\lambda_{abs}$, wavelength of maximal absorption; $\lambda_{em}$, wavelength of maximal emission; $\varepsilon$, molar absorptivity at $\lambda_{abs}$ (standard error is typically 10%); $\phi$, fluorescence quantum yield; $K_D$ thermodynamic dissociation constant.

| Tag | Fluorogen | $\lambda_{abs}$ (nm) | $\lambda_{em}$ (nm) | $\varepsilon$ ($mM^{-1}cm^{-1}$) | $\phi$ | $K_D$ (μM) | Ref. |
|---|---|---|---|---|---|---|---|
| frFAST | HPAR-3OM | 555 | 670 | 45 | 0.21 | 1.0 | this study |
| | HMBR | 484 | 550 | 56 | 0.13 | 0.50 | this study |
| | HBR-3,5DM | 500 | 565 | 55 | 0.50 | 0.80 | this study |
| | HBR-3OM | 498 | 566 | 55 | 0.15 | 0.80 | this study |
| | HBR-3,5-DOM | 525 | 600 | 51 | 0.19 | 3.9 | this study |
| FAST | HMBR | 481 | 540 | 45 | 0.23 | 0.13 | 1 |
| | HBR-3,5DM | 499 | 562 | 48 | 0.49 | 0.08 | 1 |
| | HBR-3OM | 494 | 561 | 40 | 0.36 | 0.31 | 1 |
| | HBR-3,5-DOM | 518 | 600 | 39 | 0.31 | 0.97 | 1 |
| iFAST | HMBR | 480 | 541 | 41 | 0.22 | 0.07 | 1 |
| | HBR-3,5DM | 499 | 558 | 46 | 0.57 | 0.06 | 1 |
| | HBR-3OM | 495 | 560 | 39 | 0.49 | 0.20 | 1 |
| | HBR-3,5-DOM | 516 | 600 | 38 | 0.40 | 0.41 | 1 |

**Supplementary Table 2.** Properties of frFAST and various monomeric far-red fluorescent proteins. Abbreviations are as follows: $\lambda_{abs}$, wavelength of maximal absorption; $\lambda_{em}$, wavelength of maximal emission; $\varepsilon$, molar absorptivity at $\lambda_{abs}$; $\phi$, fluorescence quantum yield.

| Protein | Size (kDa) | $\lambda_{abs}$ (nm) | $\lambda_{em}$ (nm) | $\varepsilon$ ($mM^{-1}cm^{-1}$) | $\phi$ | Molecular Brightness ($\varepsilon \times \phi$) | Ref. |
|---|---|---|---|---|---|---|---|
| frFAST | 14 | 555 | 670 | 45 | 0.21 | 9,500 | This study |
| miRFP670nano | 17 | 645 | 670 | 95 | 0.11 | 10,500 | 2 |
| miRFP670 | 35 | 642 | 670 | 87 | 0.14 | 12,200 | 3 |
| miRFP703 | 35 | 674 | 703 | 91 | 0.086 | 7,800 | 3 |
| miRFP709 | 35 | 683 | 709 | 78 | 0.054 | 4,200 | 3 |
| miRFP720 | 35 | 702 | 720 | 98 | 0.061 | 6,000 | 3 |
| mIFP | 35 | 683 | 704 | 66 | 0.069 | 4,500 | 3 |

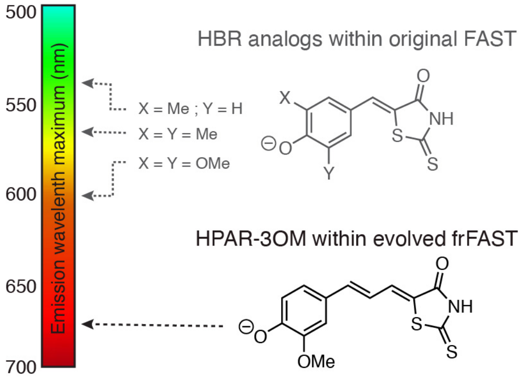


**Supplementary Figure 1.** Structures and spectral properties of HBR analogs within FAST and HPAR-3OM within frFAST.

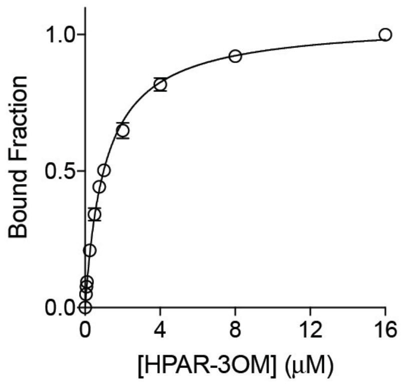


**Supplementary Figure 2. Affinity of frFAST for HPAR-3OM.** The graph shows the fraction of bound frFAST at equilibrium for various HPAR-3OM concentrations. The frFAST concentration was 0.1 μM. The titration experiments were performed at 25°C in pH 7.4 PBS (50 mM sodium phosphate, 150 mM NaCl). Data represent mean ± sem (n = 3). Least squares fit (line) gave the dissociation constant $K_D$.

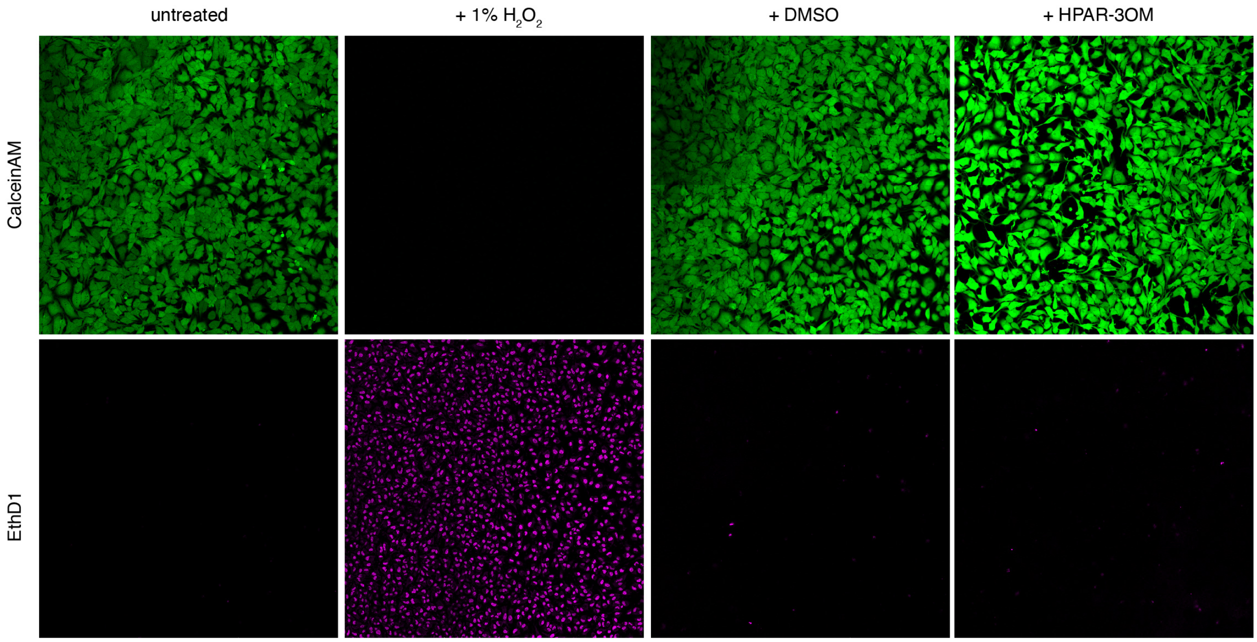


**Supplementary Figure 3. Viability assay**. HeLa cells were incubated for 24 h with solutions of HPAR-3OM at 10 µM. Cell viability was tested by using calceinAM and EthD1 (LIVE/DEAD® viability/cytotoxicity assay kit). CalceinAM is a cell-permeant profluorophore cleaved by intracellular esterases releasing the green fluorescent polyanionic calcein in live cells (green channel). EthD1 (Ethidium homodimer 1) is a non cell-permeant nucleic acid red fluorescent stain that enters only cells with damaged membranes and undergoes a fluorescence enhancement upon binding to nucleic acids, thereby producing a bright red fluorescence in dead cells (magenta channel). Control experiments with HeLa cells non-incubated with dye, incubated for 30 min with 1% hydrogen peroxide and incubated with 0.1% DMSO for 24 h are shown. Cell fluorescence was evaluated by confocal microscopy. Identical imaging settings were used for the different experiments. The experiment shows that HPAR-3OM is non-toxic for HeLa cells at the concentrations used for imaging.

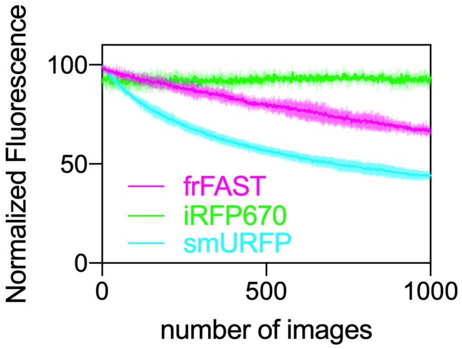


**Supplementary Figure 4. In-cell photobleaching of frFAST, iRFP670 and smURFP.** HeLa cells expressing the different far-red fluorescent reporters as fusion to H2B were imaged using a scanning confocal microscope equipped with a 633 nm laser (with a power of 21.5 kW / cm$^2$ at the specimen plane) at a frame rate of 1 image / s (pixel dwell time of 1.27 µs). frFAST was labeled with 10 µM HPAR-3OM. Traces are the mean normalized fluorescence of n = 5 cells ± SD.

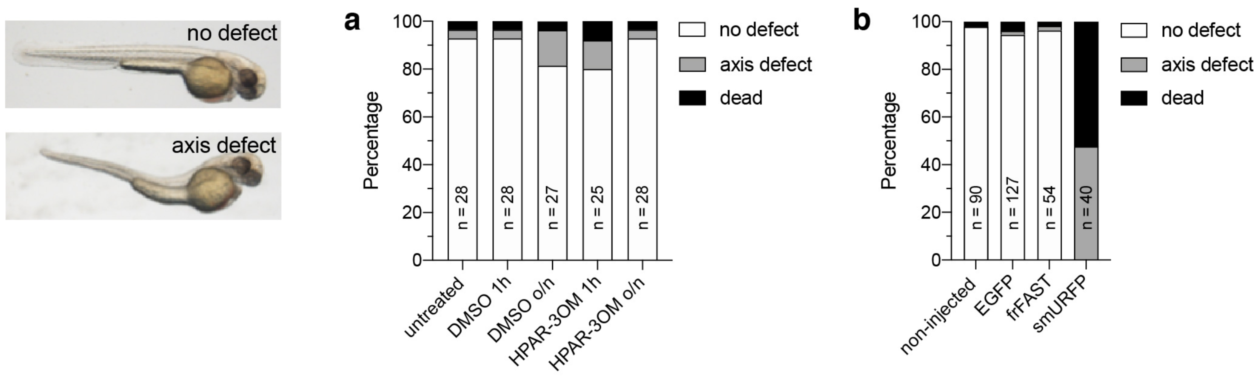


**Supplementary Figure 5. Effects of HPAR-3OM and frFAST expression on zebrafish embryos. (a)** Effect of HPAR-3OM. Zebrafish embryos were incubated with 5 μM HPAR-3OM during 1 hour at 50% epiboly or overnight (o/n) from 50% epiboly to 24 hpf. The graph shows the percentage of embryos with no defect (white), axis defects (grey), or dead (black) at 48 hpf. Controls untreated and treated with DMSO only were performed. The number (n) of embryos used for each condition is indicated. **(b)** Effect of H2B-frFAST expression. Zebrafish embryos were injected with mRNA coding for H2B-EGFP, H2B-frFAST or H2B-smURFP at the one-cell stage. The graph shows the percentage of embryos with no defect (white), axis defects (grey), or dead (black) at 24 hpf. The number (n) of embryos used for each condition is indicated.
